# Ban on wild bird importation accelerated the spread of global viral outbreaks in parrots

**DOI:** 10.1101/2022.05.31.494240

**Authors:** Ryo Komorizono, Naoyuki Aihara, Chinatsu Fujiwara, Junichi Kamiie, Keizo Tomonaga, Akiko Makino

## Abstract

Since the isolation of the first parrot bornavirus (PaBV), which causes slow-onset, fatal neurological disease in various parrot species, in the United States in 2008, PaBVs have spread rapidly worldwide; however, the reason remains unknown. In this study, we show that the most dominant lethal genotype, PaBV-4, spread via the global trade of captive birds. Analyses of traded parrot numbers in the Convention on International Trade in Endangered Species of Wild Fauna and Flora (CITES) database and PaBV-4 phylodynamics suggested that the ban of wild imported birds in the European Union (EU) in 2007 facilitated the international trade of captive parrots, which resulted in an increase in the effective population size of PaBV-4. This change coincided with a historical PaBV-4 epidemic. These data suggest that due to the low transmission efficiency and long incubation period of PaBV-4, the majority of PaBV-4 transmission occurred in breeding facilities and the increased trade of captive parrots accelerated the global spread of PaBV-4 infection. Our results indicate that interventions for the protection of wild animals and prevention of infectious diseases may conversely cause epidemics of infectious diseases in the global system. Conservation of ecosystems requires not only the establishment of importation restrictions and maintenance of the diversity of wild animals but also the implementation of multifaceted management measures, such as quarantine policies and breeding control in captive animals.

## Introduction

Birds (class *Aves*) are the most internationally traded animals (1), and the most frequently traded birds are parrots (order *Psittaciformes*), which contain a large number of threatened species according to Convention on International Trade in Endangered Species of Wild Fauna and Flora (CITES) (2). International trafficking of wild animals and plants correlates with biodiversity loss and disease transmission (3–10). In 1992, the wild bird conservation act (WBCA), which limited or prohibited the importation of exotic bird species, was enacted in the United States for the conservation of wild birds (11), and in 2007, the European Union (EU) banned the importation of wild birds to prevent invasion of the avian influenza A virus H5N1 subtype (12). However, little international trade of captive birds has been restricted, and its impact on disease transmission is completely unknown.

Avian bornavirus (ABV) infection is an emerging infectious disease identified in 2008 (13,14). ABV belongs to the genus *Orthobornavirus* in the family *Bornaviridae*, whose representative strain is Borna disease virus 1 (BoDV-1), which causes lethal encephalitis in horses, sheep, and humans (15–17). Parrot bornavirus 4 (PaBV-4) is the most dominant ABV genotype and causes lethal proventricular dilatation disease (PDD) in parrots worldwide (18,19). The genome of PaBV-4 comprises approximately 8.9 kb of nonsegmented single-stranded negative-sense RNA (13,14). PaBV-4 causes persistent infection, mainly in the central and peripheral nervous systems, that induces inflammation and selective loss of glial cells and neurons (20,21). The increased frequency of outbreaks of PaBV-4, with high fatality rates in various species of parrots, had caused economic damage to pet bird breeding facilities and may result in the loss of biodiversity worldwide. In contrast, outbreaks of BoDV-1 have been decreasing and occur in only limited habitats of its reservoir, shrews (16). On the other hand, the only evidence of PaBV-4 infection in wild birds is that viral RNA was detected in a low percentage of feces from migratory birds in Japan, but to date, there is no evidence that PaBV-4 is spreading in wild birds other than parrots (22). Contact transmission of PaBV-4 was inefficient in cockatiels (23), suggesting that even if a few wild birds are infected with PaBV-4, wild birds with low virus loads cannot be the source of viral infection in captive birds. Therefore, how PaBV-4 has spread across continents despite its low transmission efficiency remains to be elucidated.

Virus phylodynamic analysis by a Markov chain Monte Carlo (MCMC) inference method is useful for the assessment of virus transmission (24–27). It clarifies the viral dispersal history without sampling bias through the estimation of a viral common ancestor lineage and the time to most recent common ancestor (MRCA) by using data on viral genomic sequences and that date and location of virus isolation (28,29). Fluctuation analyses of effective population sizes through phylodynamic analysis can estimate the presence of viral selective pressures and circulation patterns and evaluate whether virus control strategies, such as vaccination, restriction of host trafficking, or vector control, are effective (26,27,30).

Here, to elucidate the transmission dynamics of PaBV-4, which has spread across continents despite low transmission efficiency, we investigated an outbreak of PaBV-4 in Japan and performed phylodynamic analyses of all strains reported from 2008 to 2018. Our study demonstrates that the recent global spread of PaBV-4 may be related to the increased international trade of captive birds due to the ban on wild bird imports. This finding raises the issue of the trade in rare animals in the control of infectious disease.

## Results

### High prevalence of PaBV-4 in pet bird breeding facilities

From December 2017 to February 2018, ten captive birds died in a pet bird breeding facility in Japan. This facility mainly bred imported captive parrots. The dead birds presented PDD (Fig. 1A), and plexal nonsuppurative peripheral neuritis in their gizzards was observed on pathological examination (Fig. 1B). Since an outbreak of ABV was suspected, we performed immunohistochemical (IHC) staining using an antibody against bornavirus P protein and detected viral antigens in the gizzards of all the dead birds (Fig. 1C). RT–PCR assays revealed that the ABV genotype in the brains of the dead blue-and-yellow macaws (*Ara ararauna*) was PaBV-4 (Fig. 1D). PaBV-4 was isolated from three available brain homogenates that were positive on IHC staining and RT–PCR (Table 1); these strains were named AR18A, AR18B, and AR18C, respectively (Fig. 1EF). We previously established a sensitive reverse–transcription loop–mediated isothermal amplification (RT–LAMP) assay for the detection of PaBV-4 (31). Using the assay, we assessed the PaBV-4 prevalence in feces from 25 birds with anorexia and 3 healthy birds in the breeding facility. PaBV-4 RNA was detected in half of the samples (14/28), including one from a healthy bird (Table 1). These results indicated that PaBV-4 was highly prevalent and caused gastrointestinal disorder, feather picking disease, and PDD in parrots in the facility.

**Table 1.** Summary of PaBV-4 epidemiology in the pet bird breeding facility.

|  | Species | Common name | RT-LAMP | RT-PCR | IHC | Export country | Conditions | Sequence |
| --- | --- | --- | --- | --- | --- | --- | --- | --- |
| Dead birds | A *Ara ararauna* | Blue-and-yellow macaw | + | + | + | Singapore | PDD, Peripheral neuritis | Strain AR18A |
|  | B *Ara ararauna* | Blue-and-yellow macaw | + | + | + | Singapore | Peripheral neuritis | Strain AR18B |
|  | C *Ara ararauna* | Blue-and-yellow macaw | + | + | + | Singapore | Peripheral neuritis | Strain AR18C |
| Tested birds | 1 *Cacatua alba* | Umbrella cockatoo, White cockatoo | + | + | N.T.* | Belgium | Anorexia, Weight loss | Completely identity with AR18A |
|  | 2 *Ara severus* | Chestnut-fronted macaw | - | - | N.T. | Philippines | Ataxia |  |
|  | 3 *Eolophus roseicapilla* | Rose-breasted cockatoo, Galah | + | + | N.T. | Belgium | Convulsion | Completely identity with AR18A |
|  | 4 *Poicephalus robustus* | Cape parrot | - | - | N.T. | Singapore | Healthy |  |
|  | 5 *Eolophus roseicapilla* | Rose-breasted cockatoo, Galah | - | - | N.T. | Belgium | Convulsion |  |
|  | 6 *Poicephalus robustus* | Cape parrot | + | + | N.T. | Philippines | Anorexia, Weight loss | 29 nucleotide substitutions compared with AR18A |
|  | 7 *Eolophus roseicapilla* | Rose-breasted cockatoo, Galah | + | - | N.T. | Belgium | Bloodshot of eyes |  |
|  | 8 *Eolophus roseicapilla* | Rose-breasted cockatoo, Galah | - | - | N.T. | Belgium | Anorexia |  |
|  | 9 *Eolophus roseicapilla* | Rose-breasted cockatoo, Galah | - | - | N.T. | Belgium | Convulsion |  |
|  | 10 *Eolophus roseicapilla* | Rose-breasted cockatoo, Galah | - | - | N.T. | Belgium | Candida |  |
|  | 11 *Eolophus roseicapilla* | Rose-breasted cockatoo, Galah | + | - | N.T. | Belgium | Candida |  |
|  | 12 *Eolophus roseicapilla* | Rose-breasted cockatoo, Galah | + | - | N.T. | Belgium | Convulsion |  |
|  | 13 *Amazona ochrocephala ochrocephala* | Yellow-crowned amazon, Yellow-crowned parrot | - | - | N.T. | Belgium | Ascaris |  |
|  | 14 *Psittacus erithacus* | Grey parrot, Congo grey parrot, African grey parrot | + | - | N.T. | Philippines | FPD**, Macrolabdosia, Bacillus |  |
|  | 15 *Forpus conspicillatus* | Spectacled parrotlet | - | - | N.T. | Philippines | Anorexia |  |
|  | 16 *Lophochroa leadbeateri* | Major mitchell's cockatoo, Leadbeater's cockatoo | - | - | N.T. | Belgium | Anorexia |  |
|  | 17 *Lophochroa leadbeateri* | Major mitchell's cockatoo, Leadbeater's cockatoo | - | - | N.T. | Belgium | Anorexia |  |
|  | 18 *Psittacus erithacus* | Grey parrot, Congo grey parrot, African grey parrot | + | + | N.T. | Philippines | Anorexia, Digestive failure | 31 nucleotide substitutions compared with AR18A: |
|  | 19 *Psittacus erithacus* | Grey parrot, Congo grey parrot, African grey parrot | + | - | N.T. | Philippines | Anorexia, Digestive failure |  |
|  | 20 *Psittacus erithacus* | Grey parrot, Congo grey parrot, African grey parrot | - | - | N.T. | Philippines | FPD, Digestive failure |  |
|  | 21 *Amazona aestiva* | Turquoise-fronted amazon, Blue-fronted amazon | - | - | N.T. | Belgium | Anorexia |  |
|  | 22 *Pionites leucogaster* | Green-thighed parrot | - | - | N.T. | Singapore | Healthy |  |
|  | 23 *Cacatua ducorpsi* | Solomons cockatoo | + | - | N.T. | Singapore | Anorexia |  |
|  | 24 *Eolophus roseicapilla* | Rose-breasted cockatoo, Galah | + | - | N.T. | Belgium | Anorexia |  |
|  | 25 *Poicephalus rueppellii* | Rüppell's parrot, Rüppell's parrot | + | - | N.T. | Singapore | Anorexia |  |
|  | 26 *Diopsittacus nobilis* | Red-shouldered macaw | - | - | N.T. | Japan | Temper |  |
|  | 27 *Aratinga jandaya* | Jandaya parakeet, Jenday conure | + | + | N.T. | Philippines | Anorexia | 37 nucleotide substitutions compared with AR18A |
|  | 28 *Diopsittacus nobilis* | Red-shouldered macaw | + | - | N.T. | Singapore | Healthy |  |

\*: Not tested, \*\*: Feather picking disease

**Figure 1.**
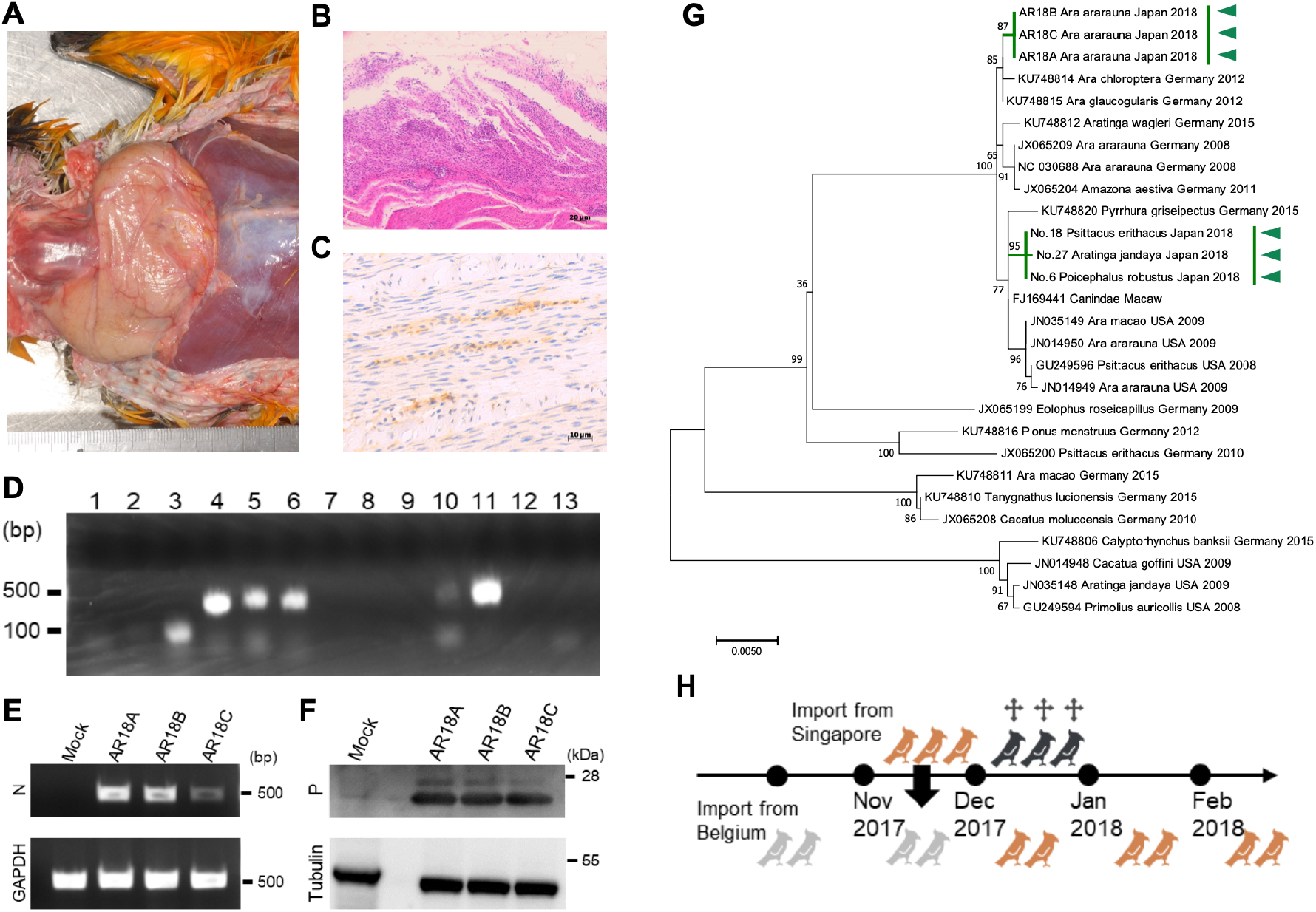
Outbreak of PaBV-4 in a pet bird breeding facility in Japan. **(A)** Proventricular dilatation in the dead blue-and-yellow macaw. **(B)** Plexal nonsuppurative peripheral neuritis in the gizzard. **(C)** IHC analysis of the gizzard. Bornavirus P antigens were stained with anti-P protein polyclonal antibody. Brown signals indicate viral antigen staining. **(D)** Viral genotype identification by RT–PCR. The primer sets are indicated as follows. 1: clade 1 specific, 2: clade 2 specific, 3: clade 3 specific, 4: universal M gene, 5: universal N gene, 6: universal N gene, 7: CnBVs, 8: ABBVs, 9: PaBV-5, 10: PaBV-2, 11: PaBV-4, 12: EsBV-1, and 13: MuBV-1. **(E)** PaBV-4 RNA detection by RT–PCR. RNA was extracted from QT6 cells inoculated with brain homogenates from three dead blue- and-yellow macaws and subjected to RT–PCR using PaBV-4 N-specific primers. GAPDH mRNA was used as the internal control. **(F)** PaBV-4 P antigen detection. QT6 cells inoculated with brain homogenates from three dead blue-and-yellow macaws were subjected to Western blotting analysis using anti-bornavirus P polyclonal antibody. Alpha-tubulin was blotted with anti-α tubulin monoclonal antibody (Sigma) as an internal control. **(G)** Phylogenetic analysis of PaBV-4 strains identified in this study. **(H)** Presumed virus transmission in the breeding facility.

Next, we determined the full genome sequences of 5 positive samples (#1, 3, 6, 18, 27) detected by RT–PCR in the tested birds. The genome sequences of sample # 1 and #3 were identical to AR18A, while those of sample #6, #18, and #27 showed 29-37 nucleotide differences compared to AR18A (Table 1). Phylogenetic analysis indicated that the AR18 strain was closely related to the German strain detected in 2012, and the strains detected in sample #6, #18, and #27 had high identity with the German strain in 2015 (Fig. 1G, indicated by green arrowheads). Sample #6, #18, and #27 were taken from birds imported from the Philippines, suggesting that these birds were infected with this strain after importation. The AR18 strain was detected in three birds imported from Singapore and two birds imported from Belgium. All Singapore birds were imported in November 2017 and died between December 2017 and January 2018. Since it takes at least 2 months for PaBVs to cause lethal infection in vivo (32–34), it is conceivable that the Singapore birds were infected with PaBV-4 before importation into Japan and that the strain was transmitted to the two birds in the breeding facility (Fig. 1H).

### PaBV-4 derived from the European strain spread worldwide

To clarify the dispersal history of PaBV-4, by using the nucleotide sequence of the PaBV-4 N gene detected from 2000 to 2018, we estimated the history of the evolutionary lineage and the common ancestor of the virus using Bayesian analysis of molecular sequences with a MCMC inference method. The maximum clade credibility (MCC) tree of PaBV-4 is shown in Fig. 2A. The Austrian strain isolated in 2000 was the closest relative to the MRCA of PaBV-4. PaBV-4 was divided into 3 evolutionary lineages: the MRCA of lineage 1 diverged in 1997 (Fig. 2A, red arrow); the MRCA of lineage 2 diverged in 1998 (Fig. 2A, blue arrow); and the MRCA of lineage 3 diverged in 1999 (Fig. 2A, orange arrow). The identified strains in this study belonged to lineage 3 (Fig. 2A, green arrowheads).

**Figure 2.**
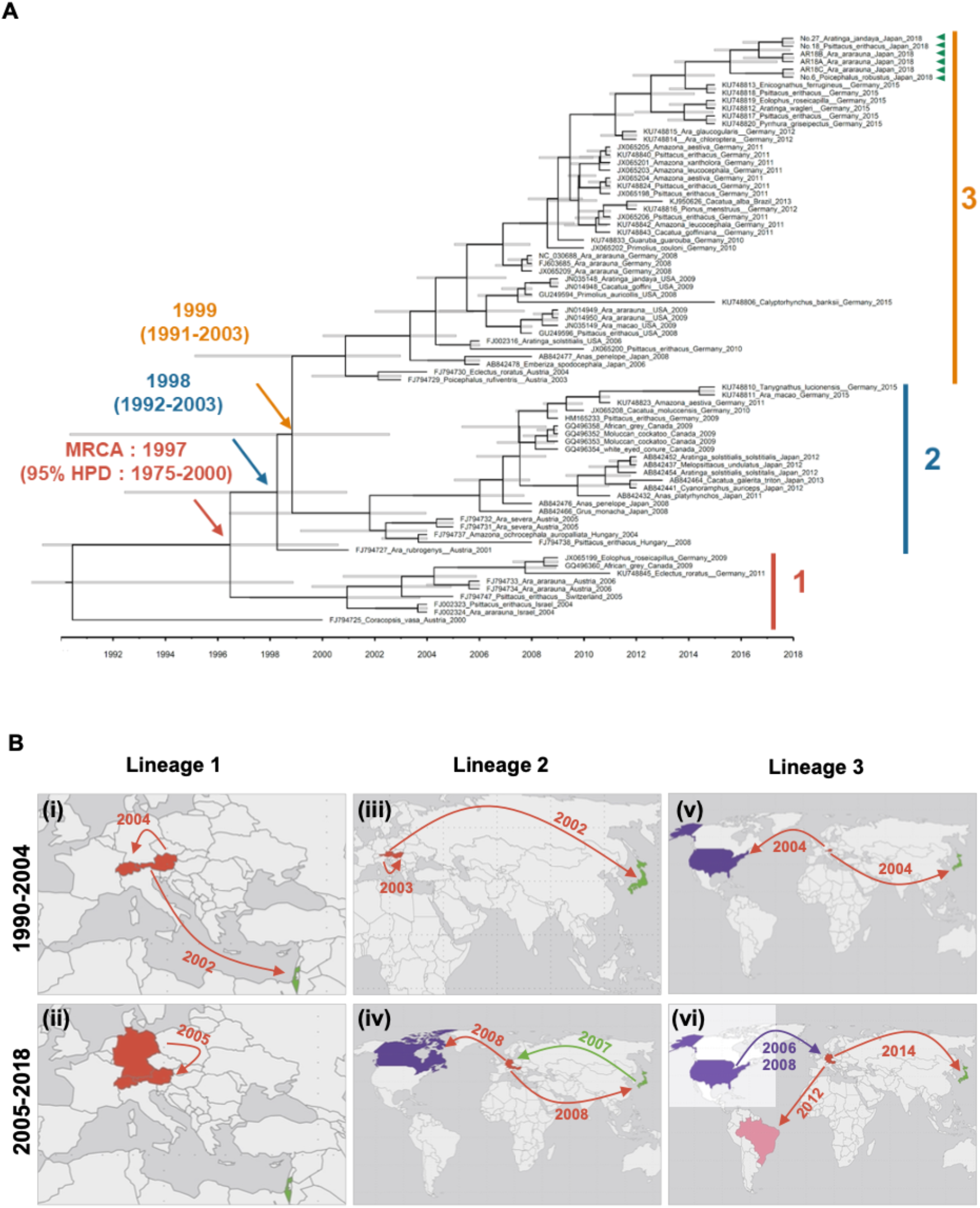
Divergence process of PaBV-4. **(A)** MCC tree of PaBV-4 from 2000 to 2018. The countries and sampling times are indicated on the tips with each accession number. The color of each node indicates the most likely location. The most recent common ancestors (MRCAs) of these strains and dates of MRCA existence are shown on the colored nodes. Mean values are shown along with the lower and upper limits of 95% highest posterior density (HPD). **(B)** PaBV-4 dispersal history from 1990 to 2018. A summary of the MCC tree is plotted on the map. (i) Lineage 1 in 1998-2004, (ii) Lineage 1 in 2005-2018, (iii) Lineage 2 in 1998-2004, (iv) Lineage 2 in 2005-2018, (v) Lineage 3 in 1998-2004, (vi) Lineage 3 in 2005-2018

The PaBV-4 dispersal history from 1990 to 2018 is mapped in Fig. 2B. In lineage 1, the Austrian strain diverged into the Israeli strain in 2002, the Swiss strain in 2004 and the German strain in 2005 (Fig. 2B i and ii). In lineage 2, the Austrian strain diverged into the Hungarian strain in 2003, and the 2002 Japanese strain evolved into the German strain in 2007 and the Canadian strain in 2008 (Fig. 2B iii and iv). The German strain derived from the Japanese strain evolved in Japan in 2008 (Fig. 2B iv). In 2004, the MRCA of lineage 3 diverged into the United States and Japanese strains (Fig. 2B v). The United Sates strain evolved into the German strain in 2006 and 2008 independently (Fig. 2B vi). The German strain that evolved in 2008 was the most dominant lineage (Fig. 2A) and diverged into the Brazilian and Japanese strains in 2012 and 2014, respectively (Fig. 2B vi). These data indicate that European-derived strains of PaBV-4 spread worldwide (indicated by red arrows in Fig. 2B) and that genetically close relevant strains were detected in geographically dispersed birds.

### The effective population size of PaBV-4 was transiently reduced after the restriction of bird trading

It was likely that PaBV-4 was transmitted worldwide via the international psittacine trade. We thus investigated the psittacine trade traffic by analyzing the CITES database from 1990 to 2017. As shown in Fig. 3A, gross international wild psittacine trading gradually decreased due to the WBCA in 1992 and the permanent ban on wild bird importation in the EU in 2007. On the other hand, gross captive psittacine trading decreased from 2000 to 2005 but continued to increase after 2006. In 2017, the gross number of captive parrots was approximately 15-fold higher than that of wild-type parrots (Fig. 3A). Next, we analyzed the fluctuation in the effective population size of PaBV-4 from 2000 to 2019 and compared it with that of BoDV-1 from 1972 to 2016, for which the number of outbreaks decreased (16). Whereas the effective population size of PaBV-4 decreased after 2007 and gradually recovered (Fig. 3B), that of BoDV-1 continued to decline (Fig. 3C). These results suggest that although psittacine trade restriction transiently reduced PaBV-4 transmission, the increase in captive psittacine trading repromoted virus transmission.

**Figure 3.**
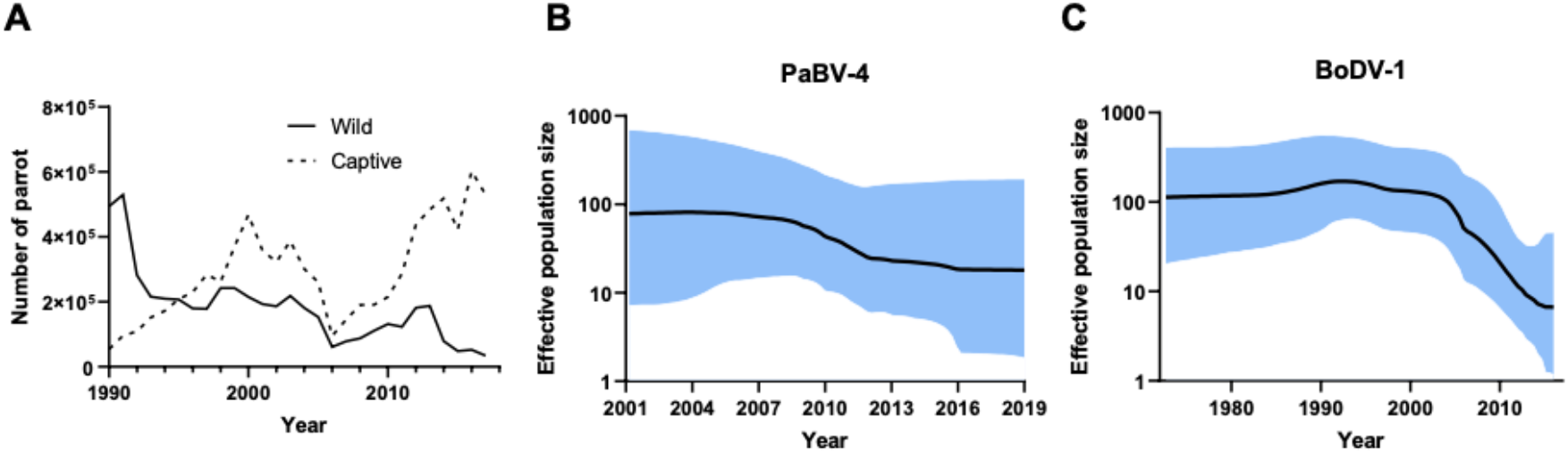
Fluctuation analysis of effective population size and the gross trade of wild and captive parrots. **(A)** The number of wild or captive parrots traded. **(B)** Effective population size of PaBV-4. **(C)** Effective population size of BoDV-1.

### The international trade network of captive parrots was consistent with the PaBV-4 dispersal history

To evaluate the involvement of the international psittacine trade in PaBV-4 transmission, we analyzed fluctuations in gross net imports and exports of wild and captive parrots using the CITES database. From 1990 to 2004, Europe mainly imported wild parrots, and Africa mostly exported them (Fig. 4A); however, after the ban on wild bird importation in the EU in 2005, North America and Asia were the main importers, and South America was the main exporter (Fig. 4B). As shown in Fig. 3A, the gross captive psittacine trade exceeded the gross wild bird trade since 1995 and continued to increase. Europe mainly imported captive parrots as opposed to wild parrots from 1990 to 2004 (Fig. 4C); however, Asia was the main importer, and Africa and Europe were the main exporters from 2005 to 2017 (Fig. 4D). In particular, the trade patterns of captive parrots exported from Europe to Asia, North America, and South America after the restriction of wild bird trade (Fig. 4D) were consistent with the dispersal history of PaBV-4 (Fig. 2B). Since PaBV-4 caused an outbreak after the ban on wild bird importation in 2005 in Europe, it is likely that the viruses that invaded before 2005 became endemic in captive birds and were transmitted worldwide via international trade of captive parrots.

**Figure 4.**
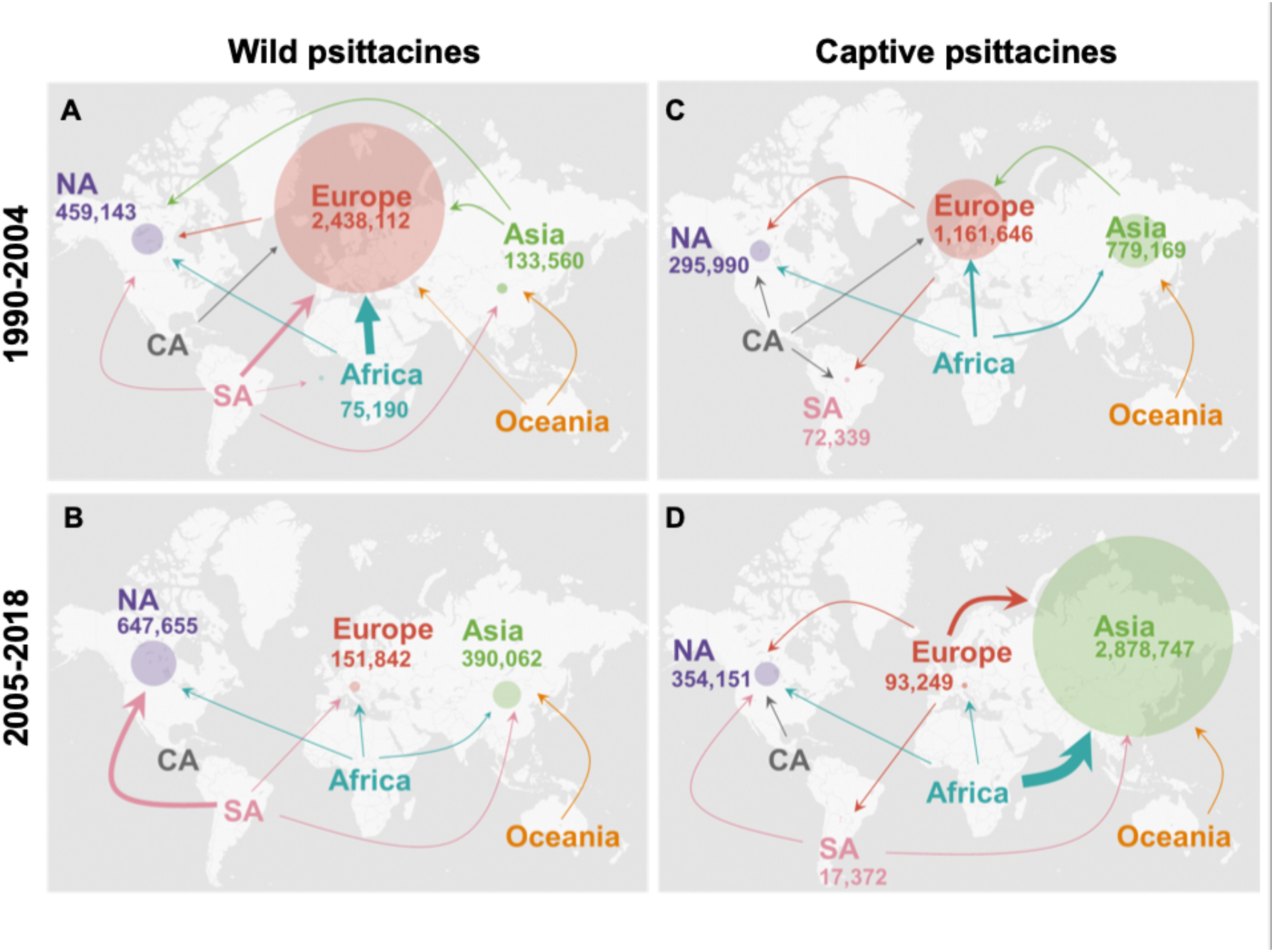
International trade network of wild and captive parrots from 1990 to 2018. International trade network of (A) wild parrots in 1990-2004, (B) those in 2005-2018, (C) captive parrots in 1990-2004, and (D) those in 2005-2019.

## Discussion

In this study, through the investigation of virus outbreaks in breeding facilities in Japan, we revealed that PaBV-4 became highly prevalent once the virus established in closed facilities. It takes 1–2 months to develop symptoms after PaBV infection (32–34), resulting in the silent spread of the virus in facilities. Since parrot breeding is expensive and time consuming, infected birds in facilities are isolated but not euthanized. The features of the virus together with captive breeding protocols may be the reason for the high prevalence of PaBV-4 in parrot breeding facilities.

An analysis of the PaBV-4 effective population size combined with CITES data showed that the ban on wild bird importation in the EU in 2007 temporarily decreased the PaBV-4 population but increased the trade of captive birds, which promoted virus transmission (Fig. 3). Furthermore, after the wild bird trade restrictions, the international trade pattern of captive parrots was consistent with the PaBV-4 dispersal history from Europe to North America, South America, and finally Asia (Fig. 2B and Fig. 4CD). Due to the low transmission efficiency and long incubation period of PaBV-4, it is highly likely that the silent spread of this virus within breeding facilities in various countries resulted in the international epidemic of PaBV-4 due to the trade of captive parrots. For instance, Asia imported a substantial number of captive parrots from 2005 to 2017. Thus, the region likely became and will likely remain a PaBV-4 hot spot. Indeed, the first report of a high prevalence of PaBV-4 in captive parrots with PDD-like disease in Thailand was published in 2019 (35). The Thailand strain is a close relative to the most dominant lineage, which is derived from the German strain (35). These data support our speculation that PaBV-4 invaded southeastern Asia via the captive parrot trade. These observations reveal that arrangements meant to prevent infectious diseases and protect wildlife can, at the same time, cause outbreaks of infectious diseases within the global trade system. There is a need for further investigation in quarantined birds to determine whether the transmission of PaBV-4 is actually due to the trade of parrots.

On the other hand, the extent of the spread of PaBVs by wild parrots remains largely unknown. Wild parrots are mainly found in tropical or subtropical regions in the Southern Hemisphere, but there have been few reports of PaBV-4 epidemics in these areas. One report described that PaBV-4 RNA was detected in the tissues of three captive-bred macaws with PDD originating from the same breeding facility in South Africa (36), implying that in Africa, transmission of the virus also mostly occurs in breeding facilities. Nevertheless, the viral sequence information of the African strain was unavailable, so we could not phylogenetically analyze the strain. Studies on the PaBV-4 prevalence in wild parrots will contribute to the elucidation of further details of virus transmission.

Recently, accidental or deliberate releases of exotic birds have resulted in establishment success in non-native habitats (3,37). In Europe, a large number of wild and captive parrots were imported as pets until 2007 (Fig. 4AC) and sometimes released into the wild; parrots accounted for 18% of established exotic birds in 2009 (37). Since these wild parrots may be a source of viral transmission, surveillance of PaBV-4 in wild parrots in Europe is important for the control of virus spread. It is feasible to control PaBV-4 transmission through the strict restriction of captive parrot trading, but this may lead to an increase in illegal and unofficial trading, which already exists (38). In other words, conservation of the ecosystem requires a multifaceted evaluation of not only importation restrictions for wild animals but also quarantine and breeding management in captive animals. PaBV-4 may be a useful model disease to study human-mediated virus transmission via international animal trade and validate the effectiveness of countermeasures, such as host trafficking restrictions and vaccination.

## Materials and Methods

### RNA extraction

Total RNA from brain tissue and cultured cells was extracted using TRIzol reagent (Thermo Fisher Scientific, Waltham, MA, USA) according to the manufacturer’s protocol. Total RNA from fecal samples was isolated using a QIAmp Viral RNA mini kit (Qiagen, Venlo, Netherlands) according to the manufacturer’s protocol or a KANEKA easy RNA extraction kit (KANEKA Corporation, Tokyo, Japan). To prepare sample solutions for the QIAmp Viral RNA mini kit analysis, fecal samples were mixed with 500 µL of RNAlater (Thermo Fisher Scientific) and centrifuged at 1,500 rpm for 5 min at 4°C. The supernatant was used for RNA extraction. For the KANEKA easy RNA extraction kit, fecal samples were mixed with 200 µL RNAlater and centrifuged at 1,500 rpm for 1 min at 4°C. After removing the supernatant, 30 µL of the extraction reagent from the kit was added to the sample, and the solution was mixed. After pipetting, 20 µL of the reagent was collected and incubated at 75°C for 5 min. This reagent was used to detect RNA in feces. RNA samples were stored in -80°C freezers before use.

### Cell culture and virus isolation

QT6 cells were cultured in Dulbecco’s modified Eagle’s medium (DMEM) Nutrient Mixture F-12 (DMEM/F12) (Thermo Fisher Scientific) containing 10% fetal bovine serum and penicillin–streptomycin. Cells were maintained at 37°C in a humidified atmosphere containing 5% CO_2_. Brain tissues from parrots were homogenized using a BioMasher V instrument (TaKaRa, Shiga, Japan) and centrifuged at 1,500 rpm for 10 min at 4°C. The homogenates were diluted with DMEM (Thermo Fisher Scientific) containing 2% FCS and centrifuged at 1,200 rpm for 5 min at 4°C. The supernatant was filtered through a 0.22 µm filter was used as the inoculum. QT6 cells were inoculated with the samples for 1 hour, and the medium was changed to DMEM/F12 containing 10% FCS, penicillin– streptomycin and amphotericin B. The inoculated cells were passaged every three or four days.

### Immunohistochemistry

Tissues were fixed with 10% buffered formalin and embedded in paraffin. An indirect immunoenzyme polymer method was used to conduct immunohistochemical analyses with anti-bornavirus P antibody in paraffined gizzard sections from dead birds. Dewaxed sections were heated in immunosaver (Nishin EM, Tokyo, Japan) at 100°C for 50 min and incubated with a rabbit anti-bornavirus P antibody diluted to a 1:1000 ratio at 4°C overnight. Peroxidase-conjugated anti-rabbit immunoglobulin G (Nichirei, Tokyo, Japan) was used as the secondary antibody. Immunoreactions were visualized using 3-amino-9-ethylcarbazole containing hydrogen peroxide (DAKO, Tokyo, Japan).

### RT–PCR

RT–PCR of extracted RNA samples was performed using universal or specific primers (shown in Supplementary Table 1). RNA was reverse transcribed using a Verso cDNA Synthesis Kit (Thermo Fisher Scientific) with random hexamers according to the manufacturer’s protocol. PCR was carried out using Tks Gflex DNA Polymerase Low DNA (TaKaRa). The PCR cycling procedure was as follows: 35 cycles at 94°C for 30 sec, 48°C for 30 sec and 72°C for 30 sec.

### Western blotting

Cells were lysed in SDS sample buffer. The cell lysate samples were boiled for 5 min and subjected to SDS–PAGE on a 5–20% gradient gel. The gel was transferred onto a polyvinylidene difluoride membrane using Trans-Blot Turbo PVDF Transfer Pack (Bio– Rad, Hercules, CA, USA). The membranes were blocked and incubated with primary antibodies against bornavirus P (1:500) and α-tubulin (Merck, Darmstadt, Germany), followed by incubation with horseradish peroxidase (HRP)-conjugated secondary antibodies (Jackson ImmunoResearch, West Grove, PA, USA). Bands were visualized with ECL Prime Western blot detection reagents (GE Healthcare, Chicago, IL, USA).

### Reverse–transcription loop–mediated isothermal amplification (RT-LAMP)

RT-LAMP assays were performed using a Loopamp RNA amplification kit (Eiken Chemical Co., Tokyo, Japan) according to the manufacturer’s protocol. Each reaction was carried out by using 40 pmol each of primers FIP and BIP, 5 pmol each of outer primers F3 and B3, and sample RNA in a 25-µL reaction volume. The reaction mixture was incubated at 63°C for 60 min, and then the reaction was terminated by heating at 95°C for 2 min. After amplification, DNA products were analyzed by agarose gel electrophoresis or visual observation using calsein FD (Eiken Chemical Co.) with a UV transilluminator or blue LED.

### Sanger sequencing

Reverse transcription was performed using SuperScript III reverse transcriptase (Thermo Fisher Scientific) with random hexamers following the manufacturer’s protocol. PCR amplification was performed by using PrimeStar MAX (TaKaRa) and 14 specific primer sets tiling the genome of PaBV-4 (shown in Supplementary Table 1). For sequencing, the PCR products were purified with a QIAquick PCR purification kit (Qiagen) and then directly sequenced with primers. PaBV-4 sequences identified in this study have been deposited in GenBank under accession numbers LC486412 to LC486417.

### Phylogenetic tree construction and population dynamic analysis

Sequences were aligned by using MUSCLE34, and phylogenetic analysis was performed by MEGA 7.0 using the maximum-likelihood (ML) methods. The ML trees were calculated using a reduced alignment excluding the outgroup sequences using a TN93 + gamma distribution model. Bootstrap support was assessed by 1,000 replications.Phylodynamic analyses were conducted on the nucleotide sequences of the N gene of PaBV-4 (162 nucleotides) from 76 isolates using the Bayesian analysis approach of molecular sequences via a Markov chain Monte Carlo (MCMC) inference method in BEAST v1.1042.43 after multiple alignments by MUSCLE. The estimates were obtained based on the Bayesian skyline tree prior to using the TN93 + gamma substitution model, the lognormal relaxed clock and empirical base frequencies. The MCMC analysis ran for 60 million iterations, with sampling every 1,000 iterations after 10% burn-in. Each effective sample size was ensured to be over 200 using Tracer v1.7. MCMC trees were created using TreeAnnotator v1.8.1 and edited in FigTree v1.4.2. A coalescent Bayesian skyline plot (BSP) was constructed by using BEAST, and mean values are shown along with the lower and upper limits of 95% highest posterior density (HPD).

### Convention on the International Trade in Endangered Species (CITES) database analyses

Data on the global importation and exportation of birds were extracted from the CITES database (https://trade.cites.org/). The database includes information about traded bird species, exporters and importers, and gross trade. Data from 1990 to 2017 were referenced using the following standards: “Live” and “Egg” for trade terms, “Wild” and “Captive” for sources, and “All purposes” for purposes. To differentiate by geographic region, trade data were classified into six regions based on the United Nations World Geographic Classification: Asia, Africa, North America, Central America, South America, and Oceania. Countries in the Middle East, such as Israel and the United Arab Emirates, were classified as “Asia”. Overseas territories, mainly in Central America and South America, such as Netherlands Antilles and French Guiana, were classified according to geography, not governing country. To accurately calculate data, duplicate data lines and records that equated importers and exporters were removed from the dataset. The number of birds in each trade event was calculated by comparing the values reported from the exporting and importing countries and adopting the larger value as the actual traded number. The number of net imports was calculated by subtracting the number of trade birds in their own region from the total amount. At that time, data on transactions for which the exporting country was “unknown” were excluded to maintain the quality of the data. Tableau and GraphPad Prism 8 were used to map the values for each region and plot the data.

## Acknowledgments

This work was supported in part by JSPS KAKENHI grant numbers JP20H05682 and JP21K19909 (KT) and JP18K05991 (AM); MEXT KAKENHI grant numbers JP16H06429, JP16K21723, JP16H06430 (all to KT), and JP19H04834 (AM); the JSPS Core-to-Core Program; AMED grant number JP19fm0208014 (KT); AMED grant number JP20wm0325011h0001 (AM); and the Joint Usage/Research Center Program on inFront, Kyoto University.

## Figures and Tables

**Table S1.**
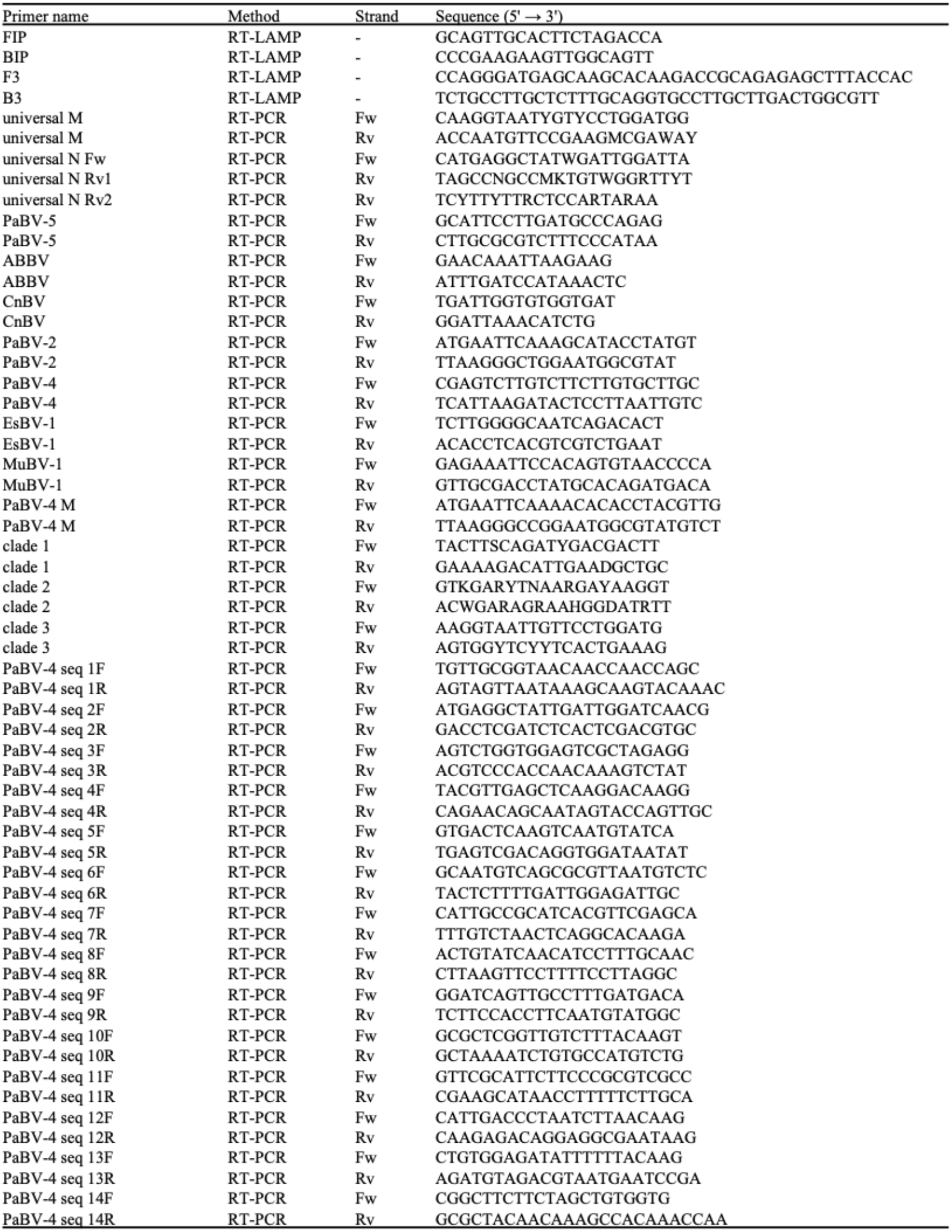
Primers used in this study.

**Table S2.** Virus sequences used in phylodynamics analysis.

| Accession number of PaBV-4 |  |  |  |  |  |  |
| --- | --- | --- | --- | --- | --- | --- |
| AB842432 | FJ002323 | FJ794737 | JN014948 | JX065204 | KU748814 | KU748842 |
| AB842437 | FJ002324 | FJ794738 | JN014949 | JX065205 | KU748815 | KU748843 |
| AB842441 | FJ603685 | FJ794747 | JN014950 | JX065206 | KU748816 | KU748845 |
| AB842452 | FJ794725 | GQ496352 | JN035148 | JX065208 | KU748817 | LC486412 |
| AB842454 | FJ794727 | GQ496353 | JN035149 | JX065209 | KU748818 | LC486413 |
| AB842464 | FJ794729 | GQ496354 | JX065198 | KJ950626 | KU748819 | LC486414 |
| AB842466 | FJ794730 | GQ496358 | JX065199 | KU748806 | KU748820 | LC486415 |
| AB842476 | FJ794731 | GQ496360 | JX065200 | KU748810 | KU748823 | LC486416 |
| AB842477 | FJ794732 | GU249594 | JX065201 | KU748811 | KU748824 | LC486417 |
| AB842478 | FJ794733 | GU249596 | JX065202 | KU748812 | KU748833 | NC_030688 |
| FJ002316 | FJ794734 | HM165233 | JX065203 | KU748813 | KU748840 |  |

| Accession number of BoDV-1 |  |  |  |  |  |  |
| --- | --- | --- | --- | --- | --- | --- |
| KY490041 | KY002071 | EU622878 | MG800681 | AY374547 | AY374537 | AY374527 |
| KY490040 | KU748787 | EU095836 | MG800680 | AY374546 | AY374536 | AY374526 |
| KY002079 | KM349819 | EU095835 | MG800679 | AY374545 | AY374535 | AY374525 |
| KY002078 | KM349818 | EU095834 | MG800678 | AY374544 | AY374534 | AY374524 |
| KY002077 | KJ127544 | DQ680833 | MG800677 | AY374543 | AY374533 | AY374523 |
| KY002076 | KJ127543 | DQ251042 | AY374552 | AY374542 | AY374532 | AY374522 |
| KY002075 | KF275185 | DQ251041 | AY374551 | AY374541 | AY374531 | AY374521 |
| KY002074 | KF275184 | DQ116030 | AY374550 | AY374540 | AY374530 | AY374520 |
| KY002073 | GQ861449 | MG800683 | AY374549 | AY374539 | AY374529 |  |
| KY002072 | EU622879 | MG800682 | AY374548 | AY374538 | AY374528 |  |

